## Supplementary Table and figures for "Molecular architecture and conservation of an immature human endogenous retrovirus"

**Supplementary Table 1 |** Data collection, subtomogram averaging and model refinement statistics of the immature HERV-K capsid.

| **Sample** | HERV-K VLP |
|---|---|
| **Data collection** | |
| Magnification | 105,000x |
| Voltage (kV) | 300 |
| Electron dose (e–/Å$^2$) | 127.5 |
| Defocus range (µm) | -2 to -5µm |
| Detector | Falcon4 |
| Energy-filter (slit) | 5eV |
| Super-res mode | N/A |
| Acquisition scheme | -60°/60°, 3°, dose-symmetric |
| Frame number | EER format |
| Pixel size (Å) | 1.503 |
| No. of Tilt-series/Micrographs | 124 |
| **Data processing** | |
| No. of Tilt-series/Micrographs | 124 |
| Symmetry imposed | C6 |
| Final particle images | 188,111 sub-volumes / 7,712,551 particle projections |
| Map resolution (Å) | 3.2 |
| FSC threshold | 0.143 |
| **Refinement** | |
| Initial model used (PDB code) | N/A |
| Model resolution (Å) | 4.0 |
| FSC threshold | 0.5 |
| Validation | |
| MolProbity score | 1.02 |
| Clashscore | 0.98 |
| Poor rotamers (%) | 0.0% |
| Ramachandran plot | |
| Favored (%) | 96.46% |
| Allowed (%) | 4.42% |
| Disallowed (%) | 0.88% |

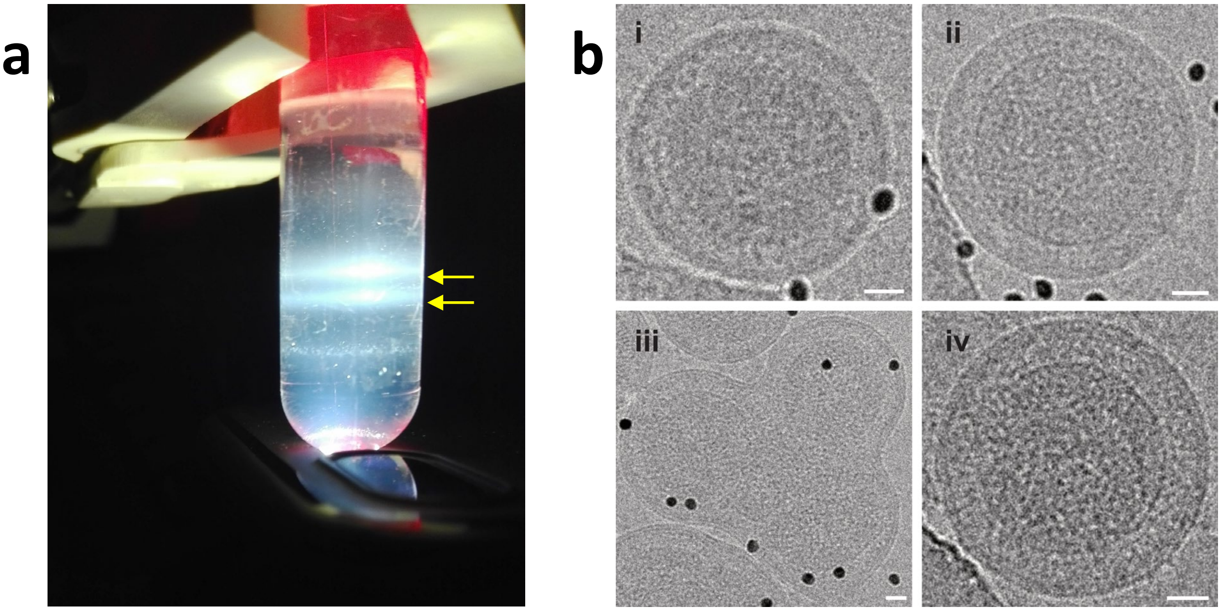

**Supplementary Figure 1 | Purification of immature HERV-K VLPs. (**a) Density gradient after ultracentrifugation. Bands visualized with a bright light. (b) Examples of VLPs in the lower band (i and ii) and upper band (iii and iv), indicated by yellow arrows. Scale bars 20 nm.

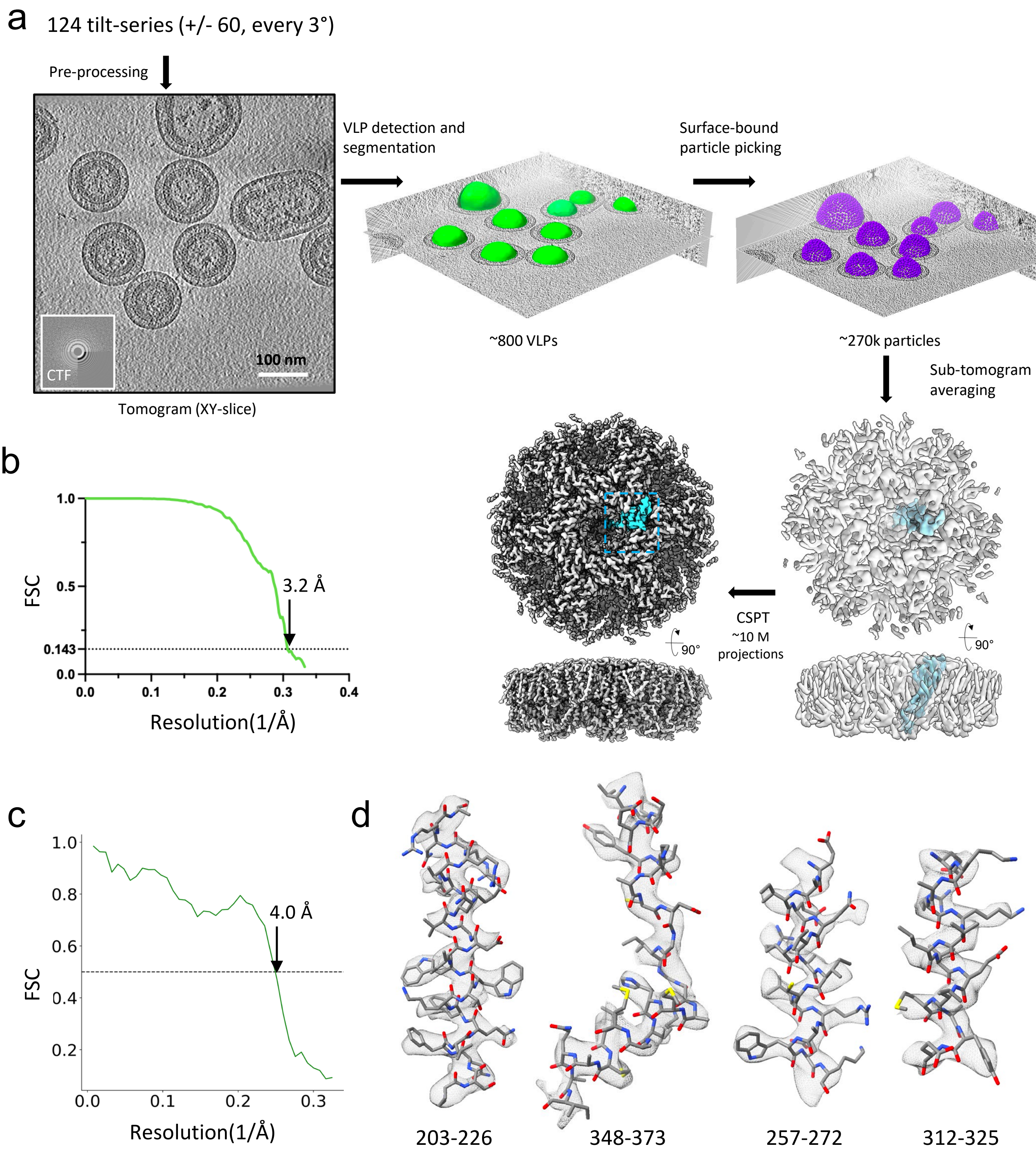

**Supplementary Figure 2 | Cryo-ET/SVA structure determination workflow.** (a) The STA flowchart illustrating steps in sub-tomogram averaging data analysis. Tomographic slice and 2D CTF profile obtained after tilt-series pre-processing. Picking of VLPs and estimated segmentations are represented as green surfaces. Automatically picked positions on the VLPs surface are represented as purple dots. Low resolution sub-tomogram average obtained from 240k particles and corresponding high-resolution map obtained using constrained single-particle tomography (CSPT) from 10M particle projections. The Gag protein monomer within the density map is colored in cyan. (b) Fourier shell correlation (FSC) curve calculated between two half-maps showing the estimated resolution using the 0.143-cutoff criteria. (c) Fourier shell correlation (FSC) curve calculated between structural model and the density map. (d) A gallery of secondary structure map features overlapped with the HERV-K immature Gag structure model.

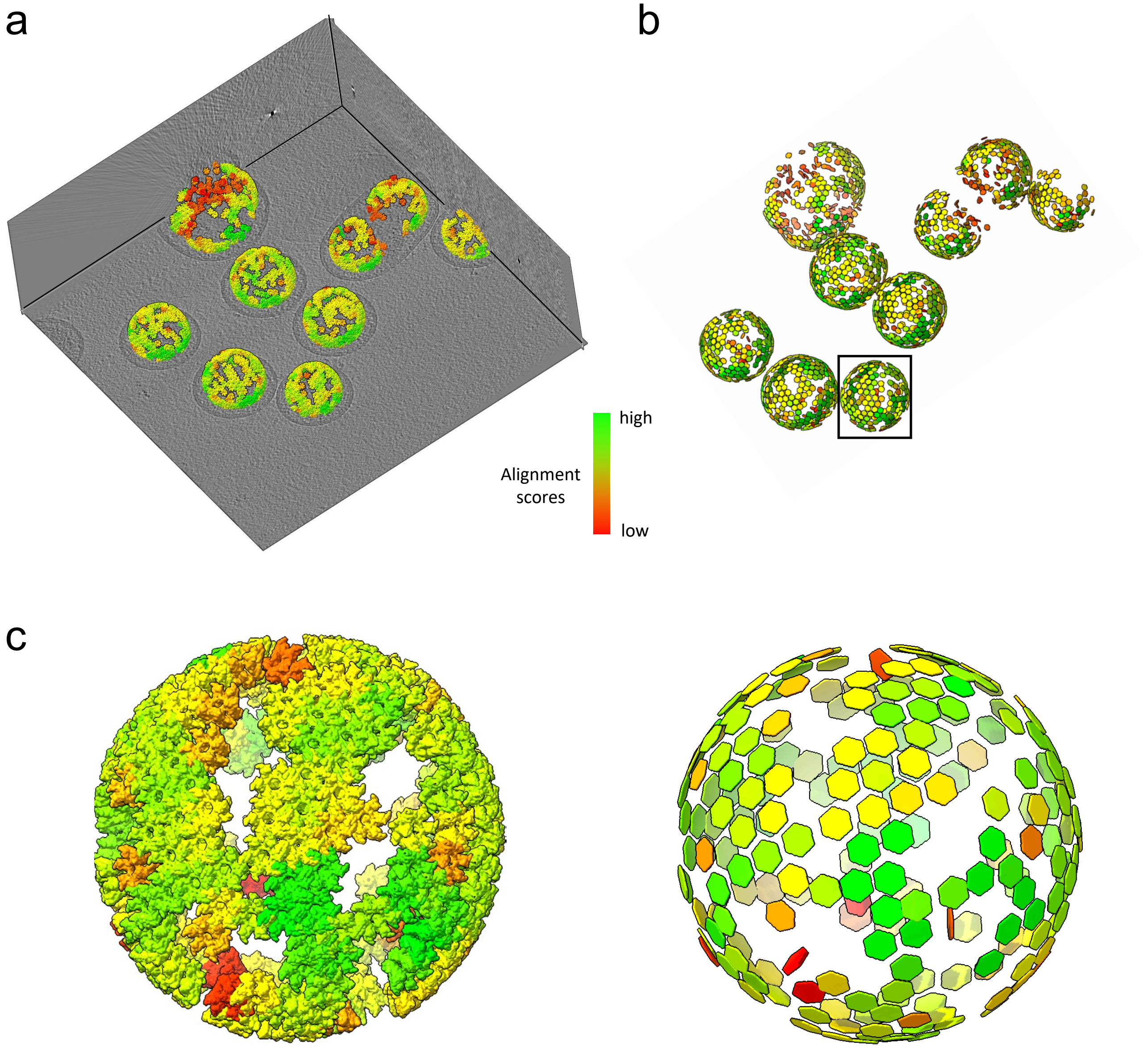

**Supplementary Figure 3 | Mapping of high-resolution models into tomograms of HERV-K VLPs.** (a-b) HERV-K hexamer units are mapped back into the original tomogram using the final alignment parameters obtained during sub-tomogram averaging. The Gag hexamer units are displayed as hexagons in 3D space and colored according to the alignment score of each particle with high values shown in green (better alignment accuracy) and low values shown in red (worse alignment accuracy). Shown in b are HERV-K VLPs extracted from the tomogram. (c) Zoomed-in view of a representative VLP (boxed in b) shows the distribution of Gag hexamers, displayed with structural models (left) or hexagons (right).

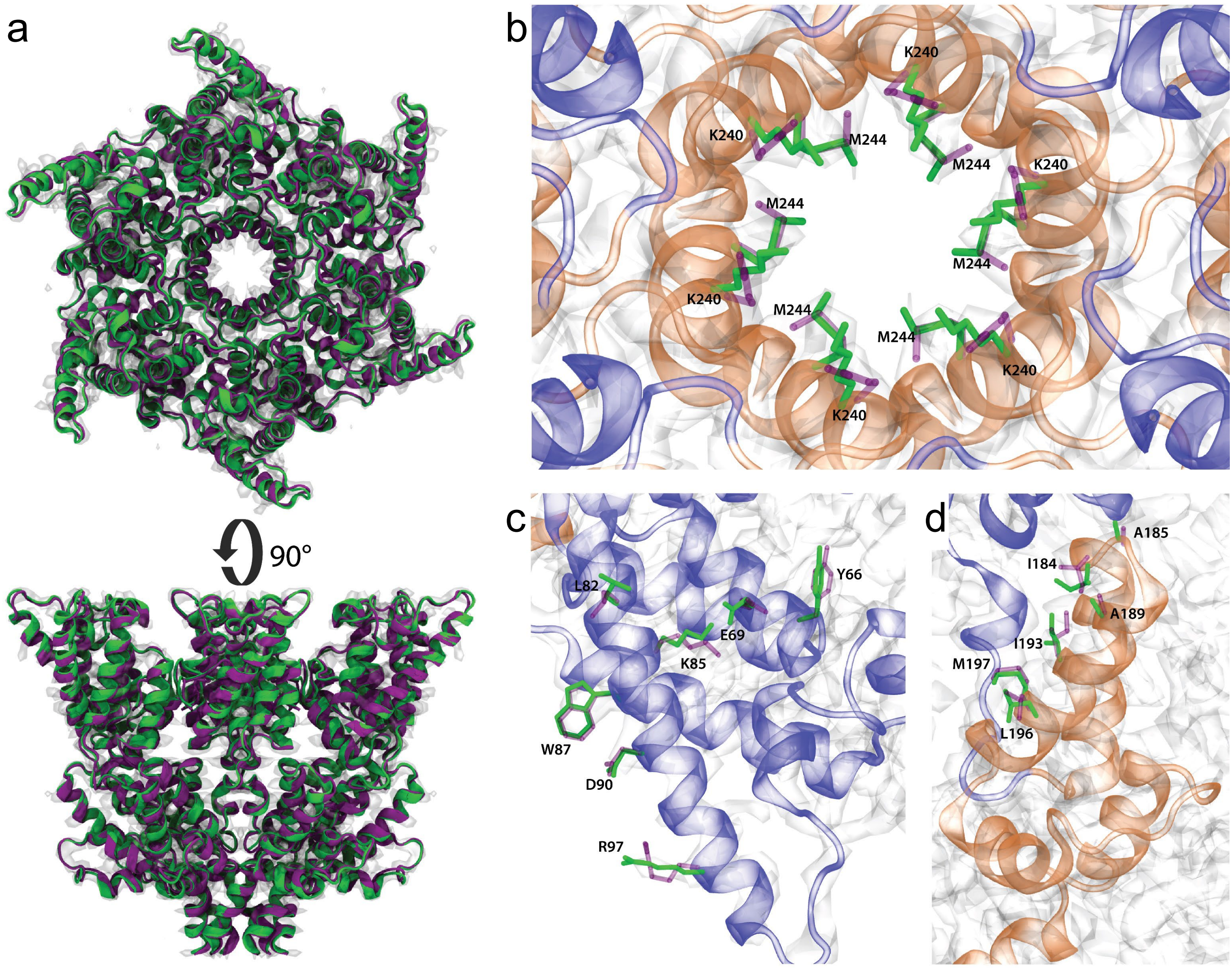

**Supplementary Figure 4 | RosettaCM refined immature HERV-K Gag structure.** (a) The original (purple) and RosettaCM refined (green) HERV-K immature CA hexamer structures. The refinement conserves the secondary structure and is consistent with the cryo-EM density (white transparent). (b-d) changes in the dihedral angles and rotamers of residues in the (b) 6HB, (c) trimer interface and (d) dimer interface due to the geometric optimization. Refined residues are shown in solid green while the original conformation is shown in transparent purple. The CTD is colored in orange and the NTD in blue.

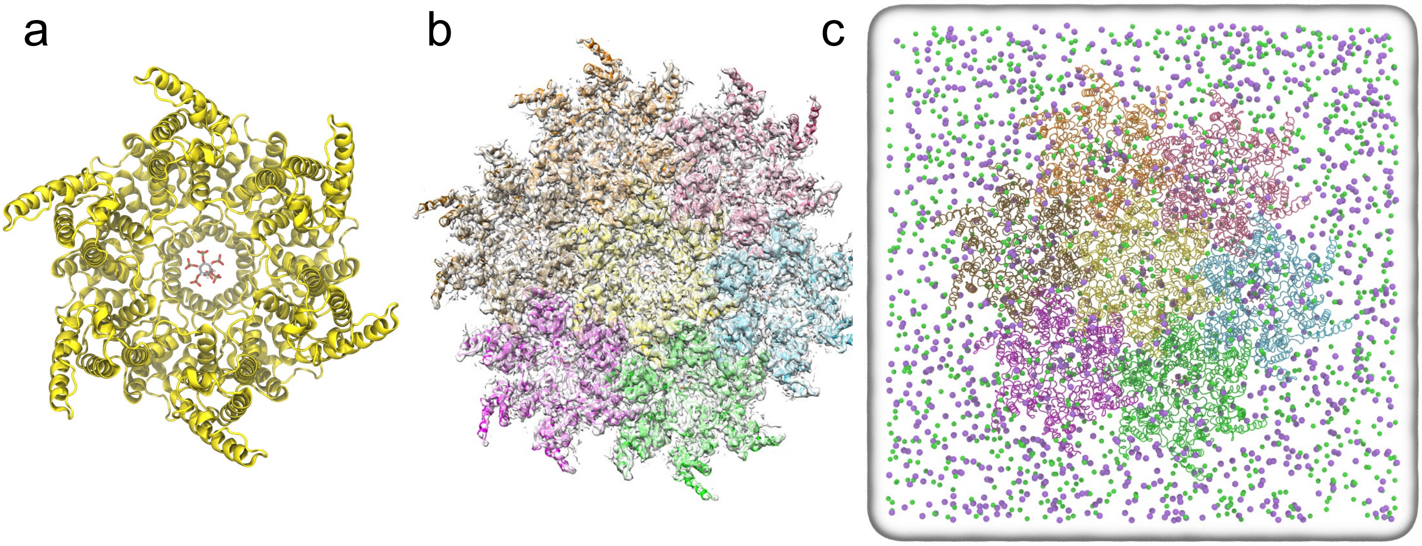

**Supplementary Figure 5 | System setup for computer simulations.** (a-b) The top-view snapshots of the immature HERV-K hexamer with a docked IP6 molecule (a) and of the HERV-K 7-hexamer lattice with IP6 molecules fitted into the cryoET STA map (white transparent volumetric density). In each snapshot, hexamers are highlighted in uniquely colored cartoon representations and IP6 molecules are highlighted in a stick representation. (c) The final simulation domain. Sodium ions are represented as purple spheres; chlorine ions as green spheres; and the water box as white volumetric density. The total number of atoms in the simulation domain is 1,247,027.

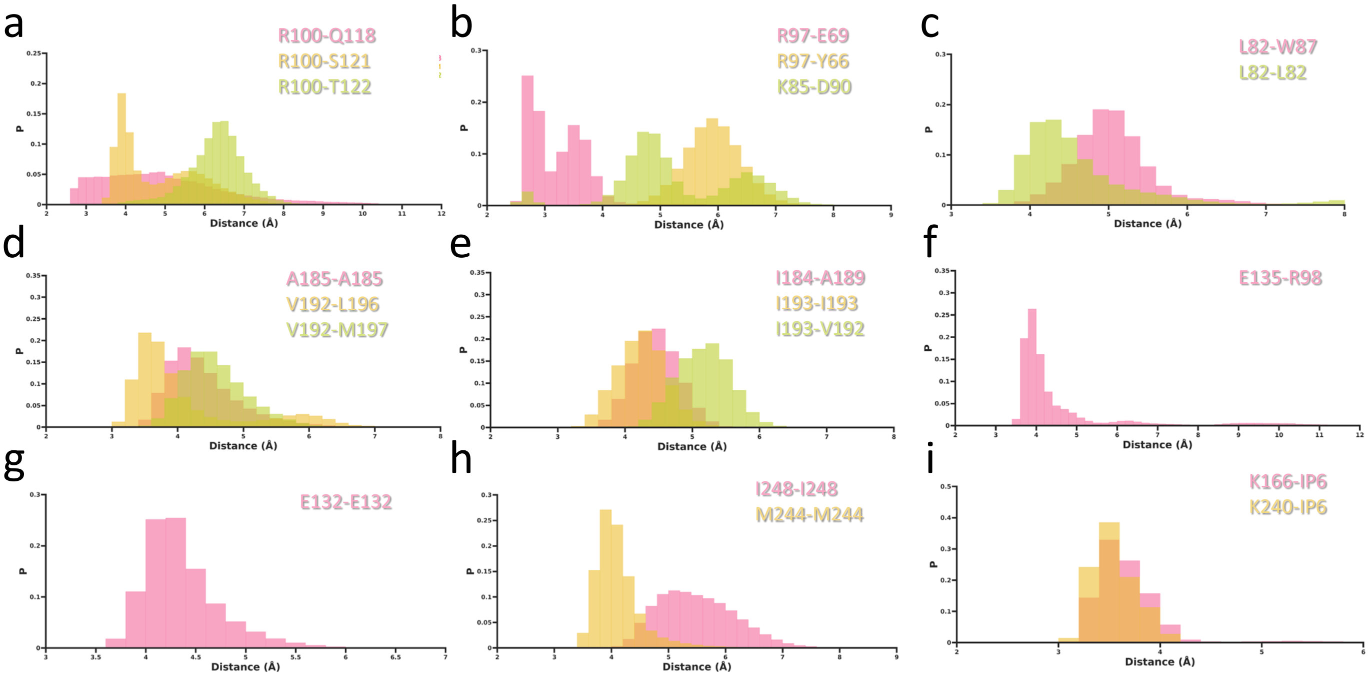

**Supplementary Figure 6 | Analysis of intermolecular interactions in computer simulations.** (a-c) The distributions of the distance metrics of the interactions at the helix 4 inter-hexamer trimer interface between R100 and Q118, S121, and T122 amino acid side chains (a), between R97 and Y66 and E69 as well as between K85 and D90 amino acid side chains (b), and between L82 and W87 amino acid side chains. (d-f) The interactions at the helix 9 inter-hexamer dimer interface between A185, V192, L197 and M197 amino acid side chains (d), between I184, A189, I193 and V192 amino acid side chains (e), and between R98 and E135 amino acid side chains. (g) The interactions between E132 amino acid side chains and a sodium ion at the helix 6 inter-hexamer dimer interface. (h) The intradomain interactions at the inter-hexamer 6HB between M244 and I248 amino acid side chains. (i) The interactions between K166 and K240 rings with the IP6 molecules.

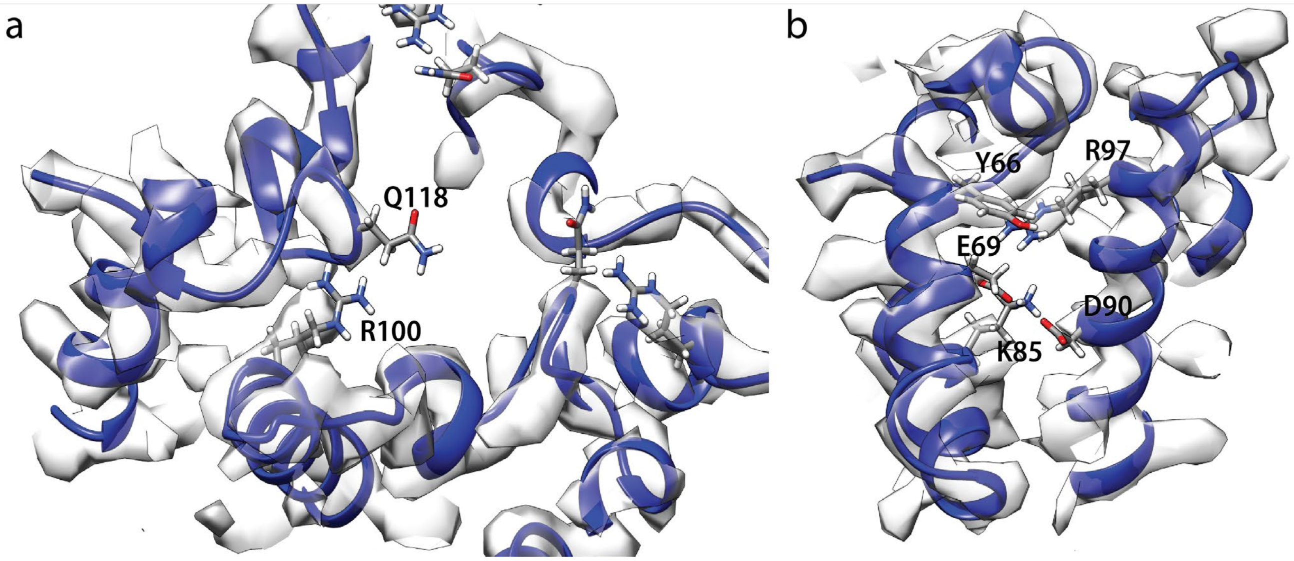

**Supplementary Figure 7 | Inter-hexamer trimer interface residue densities.** (a) Flexible residues Q118 and R100 at the inter-hexamer trimer interface present less resolved densities when compared with (b) less flexible residues Y66 and R97, as measured from MD simulations.

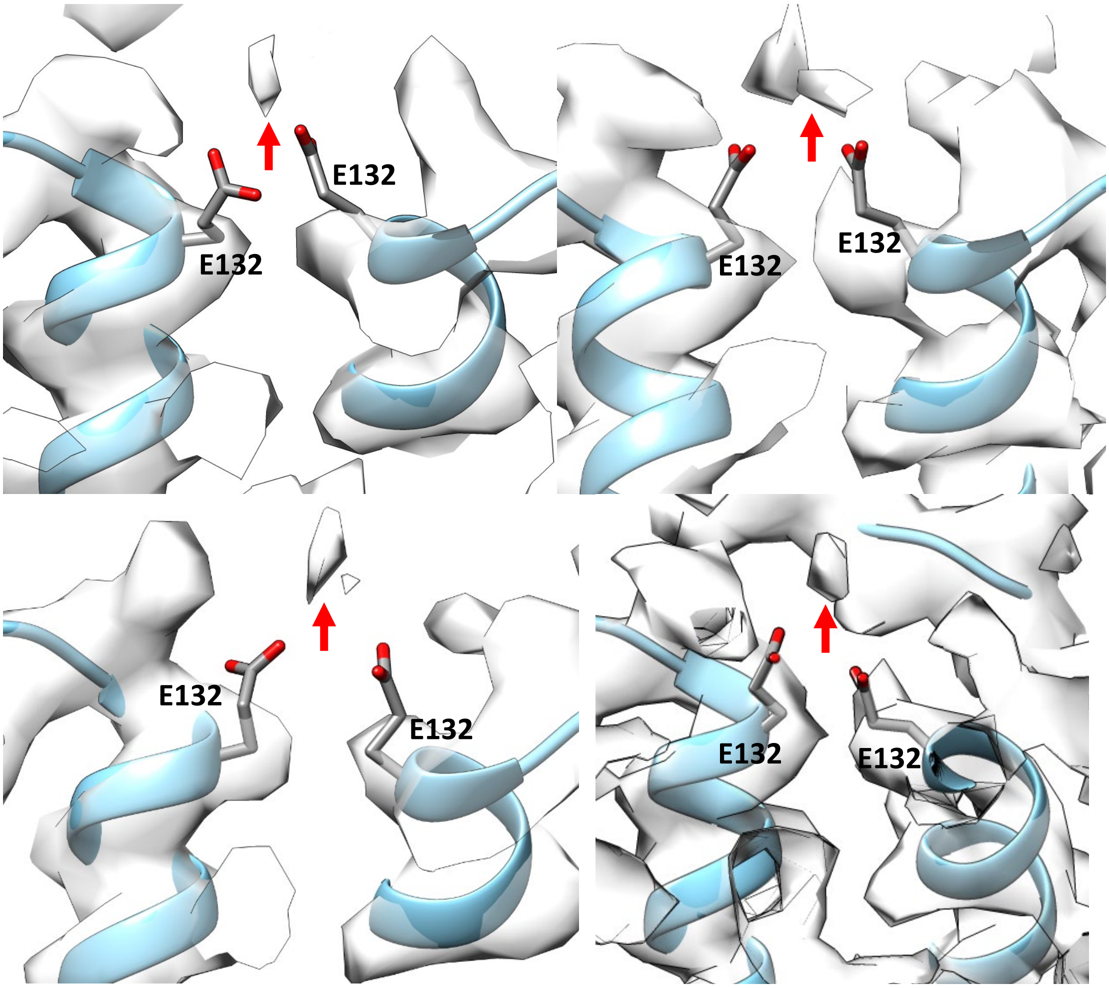

**Supplementary Figure 8 | E132 salt bridge interaction mediated by a Na+ ion.** Non protein density is observed close to the E132 sidechains at inter-hexamer interfaces. MD simulations suggest this density (red arrow) correspond to a Na+ ion coordinating ionic interactions between E132 residues in different chains.

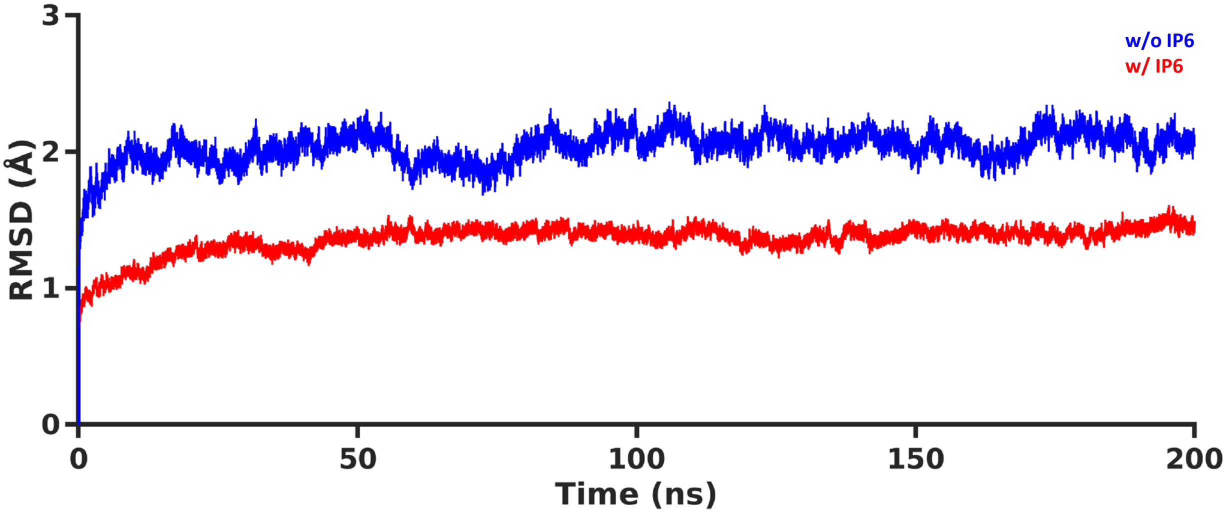

**Supplementary Figure 9 | Interactions between IP6 and two Lysine rings.** The heavy-atom RMSD time traces of the Lys166 and Lys240 rings in hexamers computed with respect to the first frame of the MD simulations in the presence (red) and absence (blue) of the IP6 molecules.

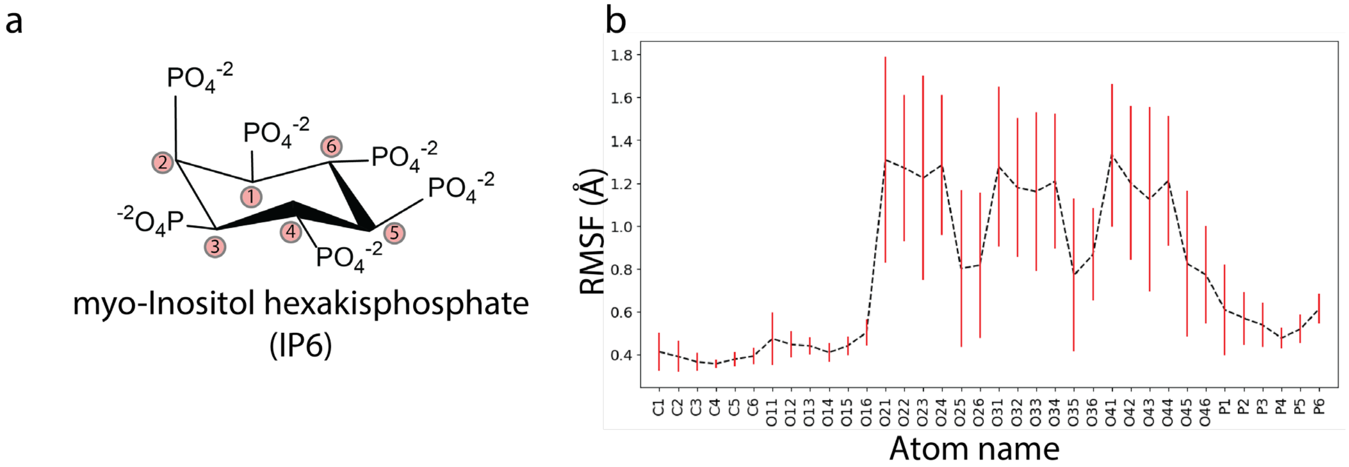

**Supplementary Figure 10 | myo-Inositol hexakisphosphate (IP6) flexibility.** (a) Chemical structure of myo-inositol hexakisphosphate (IP6). (b) RMSF of IP6 atoms averaged over the 7 IP6 molecules present in the HERV-K Gag hexamer of hexamers assembly simulated (black, dotted) and standard deviation (red). Atom names correspond to chemical element and their index in the aromatic ring.

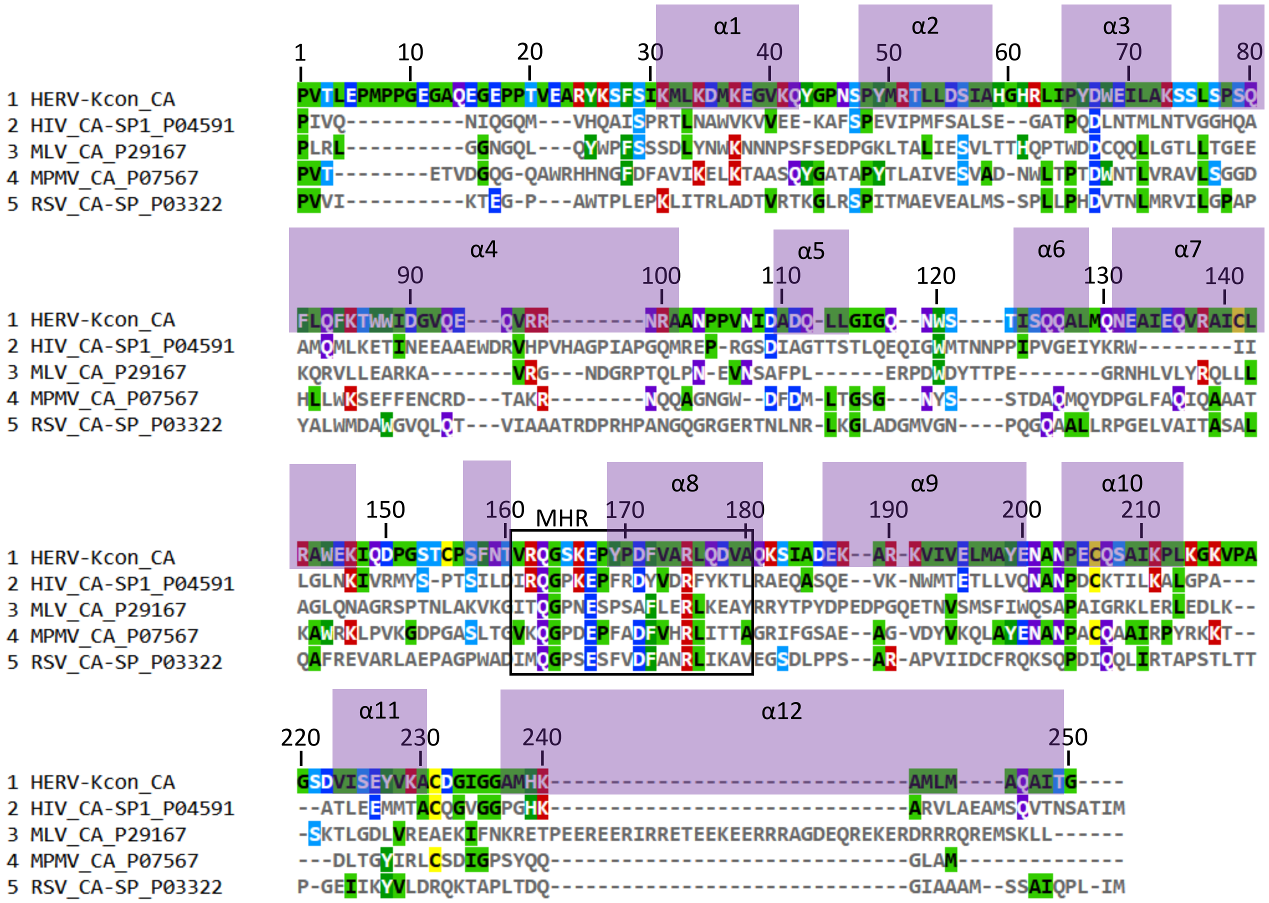

**Supplementary Figure 11 | Sequence alignment** Protein sequence alignment of CA (and the short spacer protein after CA if it exists) of the same retroviruses as in the tree in Figure 5. The helices of HERV-K$_{con}$ are labelled in purple boxes.

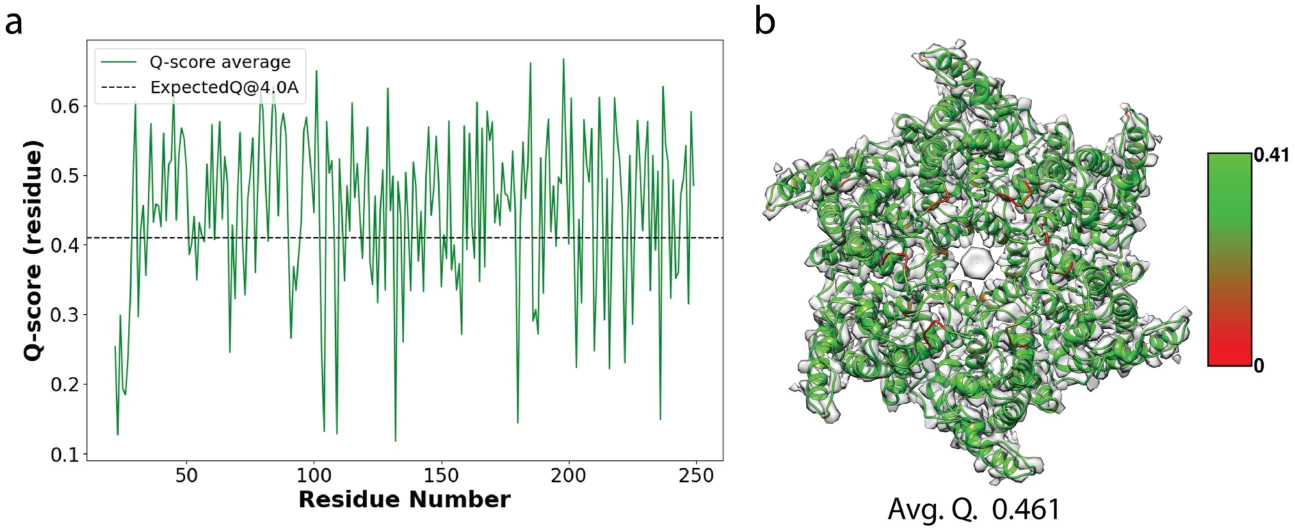

**Supplementary Figure 12 | HERV-K Gag map to model per residue Q-scores.** (a) Per residue map-to-model Q-scores for the HERV-K Gag hexamer model (green). The expected Q-score for a model of resolution 4.0 Å is 0.410 (black, dotted line). (b) The refined HERV-K Gag model colored by Q-score, good fitting is observed in helical regions while lower fitting is observed in flexible linkers. The average Q-score for the refined HERV-K Gag model is 0.461. All Q-scores have been calculated with the MapQ plugin in UCSF Chimera.
